# Acute effects of Adaptive Deep Brain Stimulation in Parkinson’s disease

**DOI:** 10.1101/749903

**Authors:** Dan Piña-Fuentes, J. Marc C. van Dijk, Jonathan C. van Zijl, Harmen R. Moes, D. L. Marinus Oterdoom, Simon Little, Peter Brown, Martijn Beudel

## Abstract

**Background:** Beta-based adaptive Deep Brain Stimulation (aDBS) is effective in Parkinson’s disease (PD), when assessed in the immediate post-implantation phase. However, the potential benefits of aDBS in patients with electrodes chronically implanted, in whom the benefits of the microlesion effect have disappeared, are yet to be assessed.

**Methods:** To determine the acute effectiveness and side-effect profile of aDBS in PD compared to conventional continuous DBS (cDBS) and no stimulation (NoStim), years after DBS implantation. 13 PD patients undergoing battery replacement were pseudo-randomised in a crossover fashion, into three conditions (NoStim, aDBS or cDBS). Patient videos were blindly evaluated using a short version of the Unified Parkinson’s Disease Rating Scale (subUPDRS) and the Speech Intelligibility Test (SIT).

**Results:** Patients had a mean disease duration of 16 years, and the mean time since DBS implantation was 6.9 years. subUPDRS scores (11 patients tested) were significantly lower both in aDBS (p=<.001), and cDBS (p=.001), when compared to NoStim. Bradykinesia subscores were significantly lower in aDBS (p=.002), but not in cDBS (p=.08), when compared to NoStim. Two patients presented re-emerging tremor during NoStim. SIT scores of patients with stimulation-induced dysarthria (11 patients tested) significantly worsened in cDBS (p=.009), but not in aDBS (p=.407), when compared to NoStim. Overall, stimulation was applied 48.8% of the time during aDBS.

**Conclusion:** Beta oscillations is effective in PD patients with bradykinetic phenotypes, delivers less stimulation than cDBS, and potentially has a more favourable speech side-effect profile. Patients with prominent tremor may require a modified adaptive control strategy.

## Introduction

Parkinson’s disease (PD) patients in advanced stages of the disease usually present with motor symptoms which are not sufficiently suppressed by dopaminergic medication, together with intolerable medication-related side-effects. Deep brain stimulation (DBS) is a successful treatment option as the disease progresses [1]. Although DBS is highly effective, both in terms of motor improvement and improvement in quality of life, there are still limitations. These include incomplete symptom suppression and side effects, such as stimulation-induced dysarthria (SID), and dyskinesias. In addition to this, most of the studies have documented the benefits of DBS only up to three years after implantation [2]. A handful of studies that have followed DBS patients in the long term (5-10 years after implantation) have suggested that the efficacy of DBS may reduce over time, especially with bradykinesia items, narrowing the therapeutic window [3,4]. Recent developments in adaptive DBS (aDBS) systems may help overcome some of these drawbacks. In aDBS, the amount of stimulation is modulated according to variations in the clinical state, or in neural activity patterns under-pinning clinical state. Here the aim is to only deliver stimulation as necessary to improve motor impairment, thereby avoiding side-effects when stimulation can be spared or lowered. Up to now, the majority of clinical studies investigating aDBS in PD have applied stimulation based on the amount of beta activity (13-35 Hz) in the subthalamic nucleus (STN) local field potential (LFP), as this biomarker is correlated with contralateral bradykinesia and rigidity [5,6], and is suppressed by DBS [7,8].

Although studies have confirmed the efficacy of ‘beta-based’ aDBS [9–13], and suggested a superior side effect profile compared to continuous, conventional, DBS (cDBS) [14,15], these have been mostly performed in newly implanted patients. Such studies are potentially compromised by the microlesion effect [16], as this confounding factor provides temporary symptom relief, which can mask the true effects of stimulation. Consequently, bradykinesia or tremor may be suppressed by the microlesion effect, and the impact of intermittent OFF stimulation periods, caused by alternately ramping stimulation up and down, thereby masked. For this reason, it is important to assess the acute effects of aDBS in chronically implanted patients. Currently it is feasible to investigate the utility of aDBS in chronically implanted patients using externalised electrodes [17], and in patients implanted with bidirectional devices [18]. Nevertheless, whether there is additional benefit over cDBS, particularly with respect to side-effects, remains unclear. One such side-effect is SID, and another is motor impulsivity, as measured by tests of response inhibition [19]. Here, we compare the clinical effect, efficiency and side-effect profile of aDBS with both cDBS and no stimulation in PD patients with long implanted electrodes in the STN.

## Materials and Methods

### Patients

We tested 13 patients with advanced idiopathic PD who were chronically treated with bilateral DBS of the STN and needed battery replacement surgery (Table 1). All patients gave written informed consent to the study protocol, which was approved by the local ethics committee. The study was registered in the Dutch Trial Register (<u>trialregister.nl</u>, trial # 5456) and the study protocol was published [20]. All patients stopped prolonged-release dopaminergic medication at least 24h prior to the measurements. In this group of advanced PD patients (mean disease duration: 16 years) the severity of OFF periods was considerably higher than in newly implanted patients. For that reason, we decided that patients skipped at least one dose of regular antiparkinsonian medication before the surgery, in order to achieve ≥6h of off-medication at the moment of the recordings. This washout period was enough to resemble the acute OFF periods that patients experience immediately after medication wears off. Battery replacement surgery was performed under local anesthesia.

**Table 1.** Clinical characteristics of the PD patients included in the study. Pt=patient, S=sex, A=age at the moment of the experiment, y=years, PD=Parkinson’s disease, M=male, F=female, LEDD=levodopa equivalent daily dose, L=left, R=right, C=case (used for monopolar stimulation). ^a^Peaks reported here were calculated offline. Therefore, they might slightly differ with the frequencies used to filter the signal intraoperatively. ^b^Due to a technical failure in one of the LFP channels, this patient was measured unilaterally in both hemispheres. ^c^This patient had the right electrode implanted in the GPi. Therefore, only results from the right hand (left electrode) were included in the analysis. ^d^This patient had the right electrode off as their standard clinical setting, as only the right hemibody required both stimulation and medication to supress clinical symptoms. However, both electrodes were tested and included in the analysis.

| Pt | S | A (y) | Time since PD start (y) | DBS use (y) | Most affected side | Tremor<sup>a</sup> | Medication | Clinical setting |  | Experimental setting |  | Freq used for filtering (±3Hz) | OFF Beta peak (Hz)<sup>b</sup> |
| --- | --- | --- | --- | --- | --- | --- | --- | --- | --- | --- | --- | --- | --- |
|  |  |  |  |  |  |  |  | Contacts used | Parameters | Contacts used | Parameters |  |  |
| PD01<sup>c</sup> | M | 58 | 24 | 9 | Left |  | Levodopa, pramipexole amantadine 1738 mg LEDD | L: -3/2+<br>R: -10/C+ | 2.9V 60µs 185Hz<br>0.8V 60µs 185Hz | L: -2/C+<br>R: -10/C+ | 1.8V 60µs 130Hz<br>1.8V 60µs 130Hz | 15 | L: 15<br>R: 15 |
| PD02 | M | 47 | 16 | 12 | Right | No | Levodopa pramipexole 1538 mg LEDD | L: -3/2+<br>R: -11/C+ | 3.5V 60µs 135Hz<br>2.2V 60µs 135Hz | L: -2/C+<br>R: -10/C+ | 2V 60µs 130Hz<br>2.2V 60µs 130Hz | 15 | L: 15<br>R: 13 |
| PD03 | M | 46 | 8 | 5 | Left | No | Levodopa duodopa 1582 mg LEDD | L: -1/2+<br>R: -11/10+ | 2.2V 90µs 180Hz<br>1.5V 90µs 180Hz | L: -2/C+<br>R: -10/C+ | 1.86V 60µs 130Hz<br>1.86V 60µs 130Hz | 20 | L: 21<br>R: 21 |
| PD04 | F | 49 | 9 | 6 | Left | Yes | Levodopa rotigotine 1558 mg LEDD | L: -1/2+<br>R: -9/10+ | 2.8V 60µs 180Hz<br>1.2V 60µs 180Hz | L: 1-/C+<br>R: -9/C+ | 2.8V 60µs 130Hz<br>1.2V 60µs 130Hz | 17 | L: 17<br>R: 19 |
| PD05 | M | 67 | 10 | 5 | Left | Yes | Levodopa 1500 mg LEDD | L: -2/C+<br>R: -11/C+ | 1.7V 60µs 180Hz<br>2.2V 60µs 180Hz | L: -2/C+<br>R: -10/C+ | 1V 60µs 130Hz<br>1.3V 60µs 130Hz | 20 | L: 22<br>R: 21 |
| PD06<sup>d</sup> | F | 64 | 23 | 9 | Left | No | Duodopa amantadine 1759 mg LEDD | L: -2/3+ | 3V 60µs 130Hz | L: -1/C+ | 2.54V 60µs 130Hz | 22 | L: 21 |
| PD07<sup>e</sup> | M | 59 | 19 | 9 | Right | Yes | Duodopa 2540 mg LEDD | L: -0/1+ | 2V 60µs 1.85Hz | L: -1/C+<br>R: -9/C+ | 2V 60µs 130Hz<br>2V 60µs 130Hz | 23 | L: 21<br>R: 21 |
| PD08 | M | 54 | 7 | 3 | Right |  | Levodopa pramipexole 725 mg LEDD | L: -0/-1/-2/3+<br>R: -9/10+ | 4.5V 90µs 190Hz<br>0.6V 60µs 190Hz | L: -1/C+<br>R: -10/C+ | 2.5V 60µs 130Hz<br>1.5V 60µs 130Hz | 15 | L: 16<br>R: 16 |
| PD09 | M | 66 | 32 | 8 | Left | No | Levodopa, pramipexole amantadine 1125 mg LEDD | L: -1/2+<br>R: -10/11+ | 4.5V 90µs 135Hz<br>4.8V 90µs 135Hz | L: -1/C+<br>R: -9/C+ | 2.05V 60µs 130Hz<br>2.05V 60µs 130Hz | 18 | L: 16<br>R: 15 |
| PD10 | M | 48 | 12 | 4 | Left | Yes | Levodopa, pramipexole amantadine 1081 mg LEDD | L: -1/C+<br>R: -9/C+ | 3.2V 120µs 130Hz<br>2.4V 120µs 130Hz | L: -1/C+<br>R: -9/C+ | 3.2V 60µs 130Hz<br>2.4V 60µs 130Hz | 29 | L: 29<br>R: 27 |
| PD11 | M | 81 | 15 | 6 | Left | No | Levodopa, pramipexole amantadine 900 mg LEDD | L: -1/2+<br>R: -10/9+ | 2.1V 60µs 135Hz<br>2.5V 60µs 135Hz | L: -1/C+<br>R: -10/C+ | 2.5V 60µs 130Hz<br>2.5V 60µs 130Hz | 17 | L: 17<br>R: 17 |
| PD12 | M | 71 | 12 | 7 | Right | No | 0 mg LEDD | L: -2/C+<br>R: -11/C+ | 3.5V 80µs 90Hz<br>3.6V 80µs 90Hz | L: -2/C+<br>R: -10/C+ | 3V 60µs 130Hz<br>3V 60µs 130Hz | 18 | L: 18<br>R: 18 |
| PD13 | M | 72 | 21 | 7 | Right | Yes | Levodopa amantadine 1098 mg LEDD | L: -2/C+<br>R: -8/C+ | 2.3V 70µs 185Hz<br>2.9V 90µs 185Hz | L: -2/C+<br>R: -10/C+ | 3V 60µs 130Hz<br>3V 60µs 130Hz | 26 | L: 26<br>R: 24 |
| Mean (±SE) |  | 60.1 (3.0) | 16 (2.0) | 6.9 (0.6) |  |  | 1318.7 (169.4) |  |  |  |  | 19.6 (1.2) | 19.2 (1.2) |

### Recording procedure

In brief, the old battery was removed and a custom-made external stimulator-amplifier device (for details see Little *et al*., 2013) was connected to the chronically implanted DBS electrodes (Figure 1). From this point the protocol took about 25-30 minutes to be completed. Bipolar STN local field potentials (LFPs) were recorded in the resting state for 30-60 seconds, using contact pairs 02 and 13. Power spectral density estimates in the beta band range and their peak frequency were calculated. The contact (either 1 or 2) in the middle of the contact pair with the greatest beta activity on each side was selected for stimulation, and the beta activity in the bipolar channel bridging the stimulation contact used for the feedback control of aDBS. A common frequency range (±3Hz) that included the beta-peak frequencies of both hemispheres, was used in each patient to filter bilateral LFPs, based on the observation that beta peak frequencies are similar between sides [21,22]. Optimal stimulation voltages were determined before the experimental conditions, based on the response to monopolar cDBS, until either optimal symptom reduction (bradykinesia and/or tremor) was achieved, side-effects occurred or voltage-dependent artefacts interfered with the recordings. Bilateral aDBS, cDBS and no stimulation (NoStim) were applied in a pseudo-randomised order, with a short washout period (2 min) between them (Figure 2). aDBS was triggered on and off when rectified beta power crossed a median power threshold up and then down, respectively.

**Figure 1.**
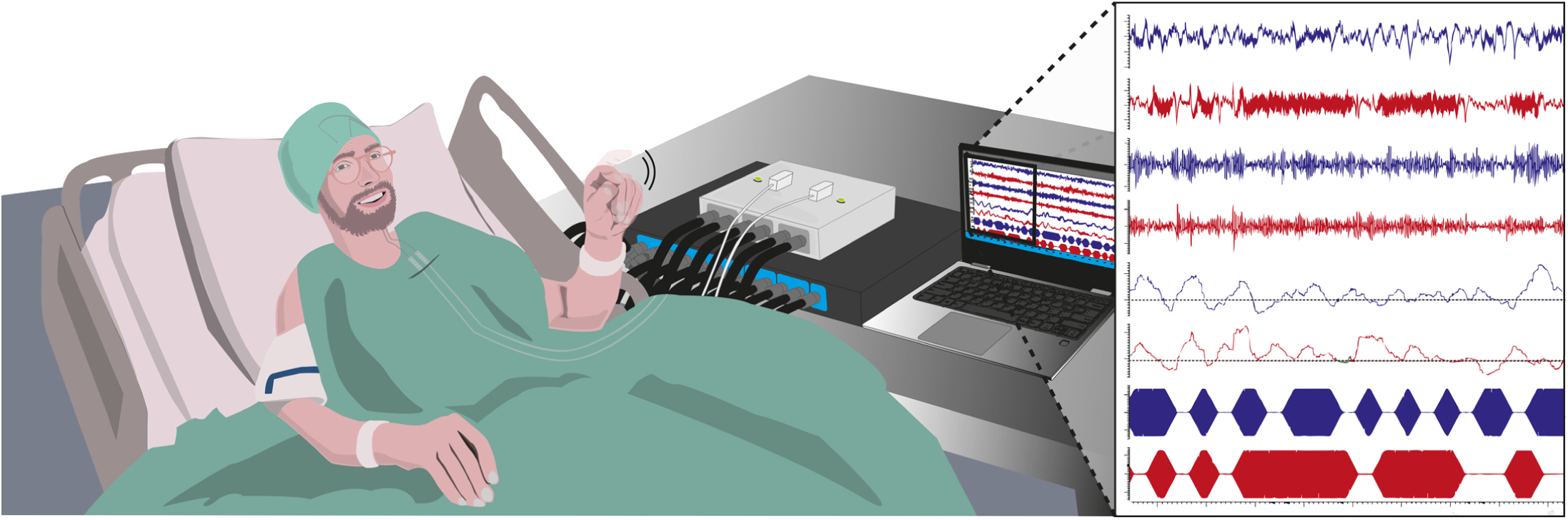
Schematic representation of aDBS. During the operation, and after the old battery is exposed, the battery is detached from the DBS electrodes and explanted. At this moment, two temporary wires are attached to the DBS electrode extension cables at the level of the chest incision, and connected to a combined stimulator and amplifier (represented here). On the right side, an aDBS recording is depicted. From top to bottom: -Original signal from left (blue) and right (red) electrodes, using a bandpass filter from 3 to 37 Hz. At this stage it is possible to appreciate artefacts caused by stimulation. -Left and right LFPs filtered around the beta peak. Here it is possible to observe the individual beta bursts. -Left and right envelopes of the beta-filtered and rectified LFPs. The dotted lines represent the thresholds selected to trigger stimulation. -Stimulation bursts, with a ramping period when stimulation is switched on and off.

**Figure 2.**
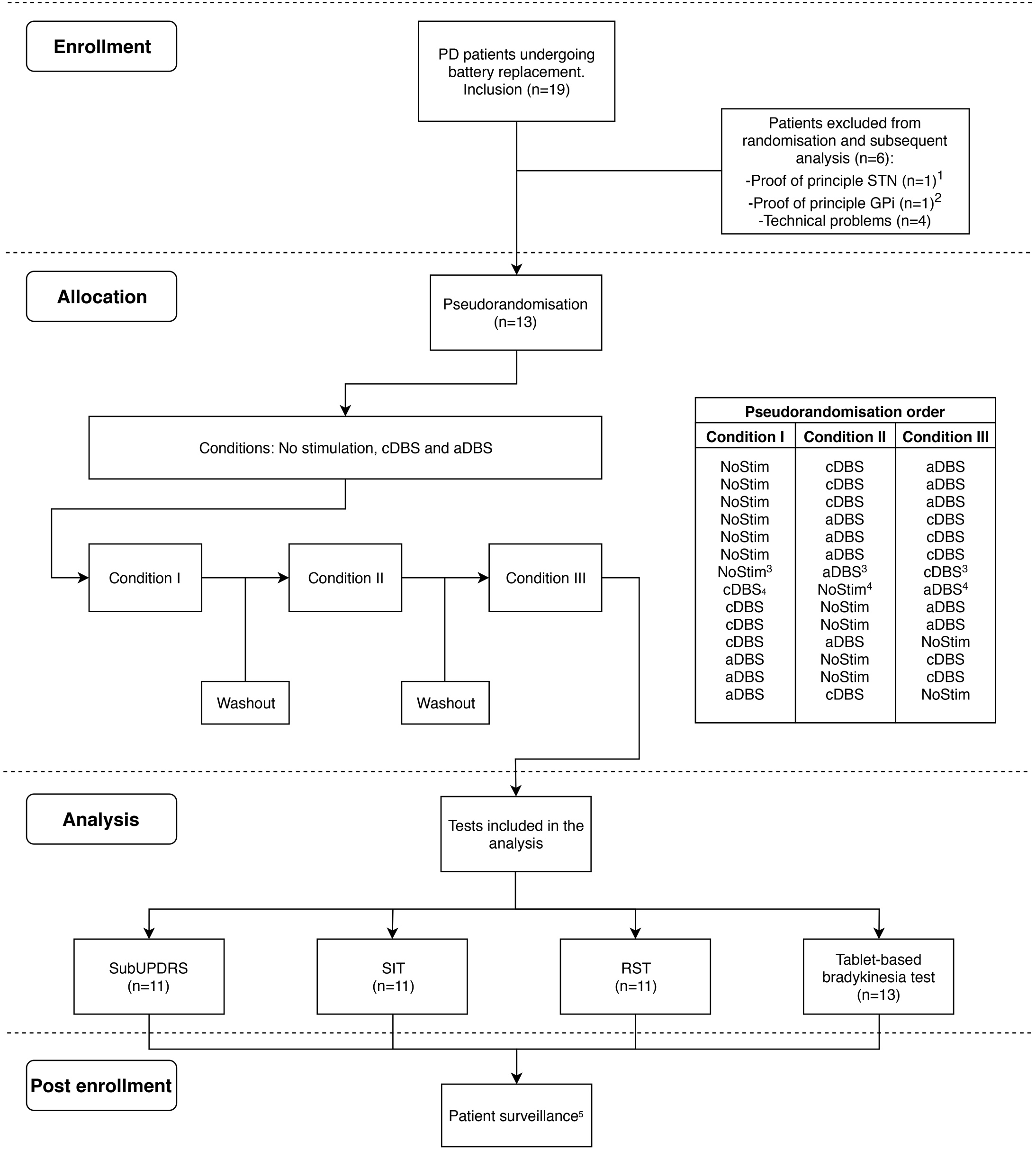
Flowchart of the inclusion algorithm and pseudorandomisation order. ^1^The protocol and tests were modified after the inclusion of this patient, who was therefore excluded. Findings have been reported in [17]. ^2^This patient was operated in the GPi, and therefore the results were reported separately in [42] ^3,4^These randomisation orders belong to the same patient (PD01). As this patient had to be measured unilaterally in both hemispheres, due to a defect in one of the aDBS box channels, each hemisphere was assigned to a different randomisation order. ^5^During patient surveillance, one patient presented a superficial infection on the surgical site, which was resolved completely after antibiotic treatment.

### Tests and data processing

During each of the three conditions, the following four tests were performed (See Pina-Fuentes *et al*., 2019 for a more detailed description of the tests):

-A short version of the motor Movement Disorders Society version of the Unified Parkinson’s Disease Rating Scale (subUPDRS) [23] was assessed and video recorded for subsequent blinded rating. Items tested bilaterally included were finger tapping, hand movements, hand pronation-supination, rest tremor and postural tremor. Videos were evaluated by three clinicians with expertise in movement disorders. Total scores were calculated for each rater/patient combination. Subgroup scores were calculated for bradykinesia and tremor items.
-The speech intelligibility test (SIT) [14], in which each participant had to read out approximately 110 words divided into 18 sentences. Sentences were recorded, and the number of intelligible words of each SIT was blindly evaluated by a speech therapist. The total amount of intelligible words of each trial was determined. This score was normalised between 0 and 1, with 0 representing completely unintelligible speech.
-Response inhibition was assessed with a tablet-based version of the reverse Stroop effect (RST) [24], with 5 congruent words and 15 incongruent words randomly presented on each trial. Reaction times were determined for each word, and the amount of correct words for congruent and incongruent conditions were calculated. Reaction times with duration ≤200 or ≥5000 ms were discarded. It has been postulated that any impaired Stroop-test responses in PD patients might be a consequence of deficits in attentional resources, rather than impairments in impulse control [25]. Accordingly we chose the RST over the Stroop test as the former arguably is less attention demanding [26].
-A bradykinesia test was performed using a tabled-based version of previously validated tapping test [27,28], in which patients had to press two separate squares in an alternating pattern (20 iterations). The first iteration of each trial was discarded, as it systematically differed from the rest, leaving 19 iterations for further analysis. Incorrect iterations were defined as each time a patient pressed the same square twice.

In the paradigm, the subUPDRS was the last test to be performed in each condition, in order to maximise the washout time between clinical scales. Hence, we use it as our benchmark measure of efficacy.

### Statistical analysis

Statistical analysis were performed using R version 3.6.0, and the statistical packages lme4 [29] and lmerTest [30]. All items were assessed for normality using Q-Q plots. Iteration times of the tapping test, and RST reaction times were log-transformed prior to further analysis. subUPDRS scores, transformed iteration times, SIT scores and transformed RST reaction times were compared using a linear mixed-effects model. Condition was treated as a fixed effect in all tests and subject as a random effect. In addition, in the subUPDRS model, randomisation order was included as a fixed effect. Additionally, bradykinesia and tremor subscores were analysed using independent models. The intra-rater absolute agreement of UPDRS-subscores was analysed using 2-way random-effect intraclass correlation coefficient (ICC). In the SIT model, the presence of SID was included a fixed effect, with SID defined as a decrement on the SIT score during cDBS, aDBS or both. In the RST analysis, congruency (congruent/incongruent) and accuracy (correct/incorrect) were added as fixed effects. In the tapping test, errors and their interaction with condition were added as fixed effects. Additionally, error counts of the tapping test and RST were analysed using a generalised linear mixed-effects model with a log-linear Poisson distribution model and an offset of 19 for each tapping-test trial, 5 for congruent words and 15 for incongruent words. In the RST error analysis, congruency was initially included as a fixed effect to test the presence of a reverse Stroop effect, and afterwards, its interaction with condition was added to the model, to test the selectivity of this effect to each condition. Models were compared using the Bayesian information criterion (BIC). The percentage time on stimulation was calculated for the aDBS condition. All results are expressed as mean ± standard error.

## Results

Patients had a mean disease duration of 16±2 years, and the mean time since DBS implantation was 6.9±0.6 years. The average levodopa equivalent daily dose used was 1318.7±169.4 mg. Of the 13 patients included in this study, all patients completed the bradykinesia test. Due to time and technical constrains, the rest of the tests were completed only by 11 patients each. All patients tolerated the test procedure, and stimulation-induced transient paresthesias were present only at stimulation voltages above those used for the experiment.

### Bilateral subUPDRS

Whereas 7/11 patients were classified as tremor dominant based on the clinical records, only five presented tremor during the recordings. Independent raters showed an excellent degree of agreement in the scores, (ICC=0.939, F(35,70)=16.312, p=<.001). The linear mixed-effect model was significant for condition, (F(2,88)=8.144, p=<.001) (Fig 3). subUPDRS scores were significantly lower in aDBS (12.30±0.85) when compared to NoStim (15.81±0.75), (t(88)=- 3.731, p=<.001), and in cDBS (12.57±0.74) when compared to NoStim, (t(88)=-3.252, p=.001), but not between aDBS and cDBS, (t(88)=-0.443, p=.659). There was no significant effect of randomisation order, (F(2,88)=2.62, p=.07). When bradykinesia subscores were analysed, aDBS scores (11.09±0.58) were significantly less than NoStim scores (12.90±1.27, t(88)=-3.096, p=.002), but cDBS scores (11.87±0.58) only showed a trend to be less than NoStim (t(88)=-1.754, p=.08). There was no difference in bradykinesia subscores between aDBS and cDBS (t(88)=1.341, p=.18). Tremor subscores were reduced with both interventions (cDBS scores, 1.33±0.73 vs NoStim 6.13±0.92, t(40)=-6.572, p=<.001 and aDBS, 2.47±0.73 vs No Stim, t(40)=-5.02, p=<.001). There was no difference in tremor subscores between aDBS and cDBS (t(88)=1.189, p=.237). However, although most patients showed tremor suppression even when stimulation was triggered off during aDBS, two patients demonstrated re-emergence of tremor when stimulation triggered off during this condition.

**Figure 3.**
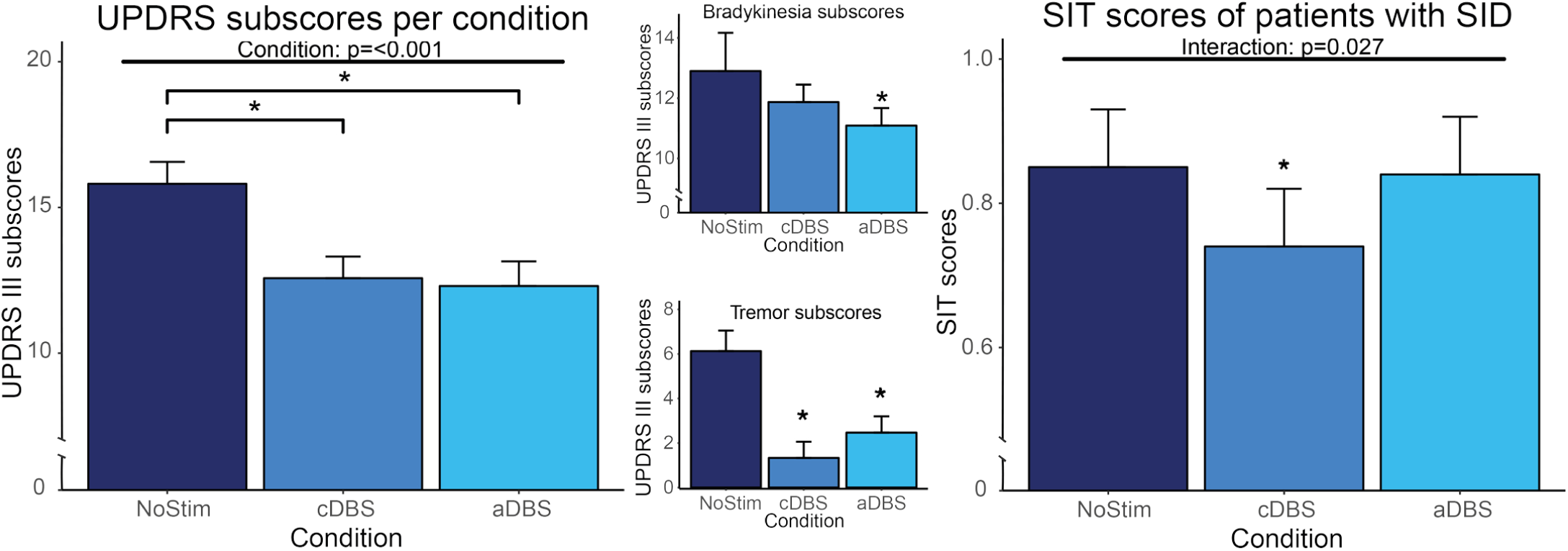
Results of the subUPDRS and SIT scores. Left: total subUPDRS scores. Upper center: subscores of bradykinesia items of the UPDRS III. Lower center: subscores of tremor items of the UPDRS III. Right: SIT scores of patients with SID.

### SIT

the linear mixed-effect model showed a significant effect for the interaction of condition and SID (F(2,22)= 4.231, p=.027). SIT scores of patients with SID significantly worsened only in cDBS (0.74±0.08) when compared to NoStim (0.85±0.08, t(22)=-2.833, p=.009), but not in aDBS (0.84±0.08) when compared to NoStim (t(22)=-0.845, p=.407).

### RST

the simpler model revealed a significant effect of congruency, (Chi^2^(1)=4.576, p=0.032), where incongruent words had a significantly increased error rate, (z=2.139, p=0.032). No significant effect of condition was found, (Chi^2^(2)= 1.772, p=0.41). A model with an interaction factor was not a better fit (BIC 146.09 vs 138.34 for the simpler model), and interaction was not significant, (Chi^2^(2)= 0.586, p=0.745). No significant effect was found in reaction times for congruency, (F(2,626.03)=0.266, p=0.766, condition, F(1, 625.99)=0.310, p=0.577), or their interaction, (F(1, 626.02)=0.723, p=0.485).

#### Tapping test

correct iteration times were reduced in cDBS (571.29±9.83ms) and aDBS (612.31±10.14ms) compared to NoStim (619.53±10.35ms). However, this reduction was only significant for cDBS, (t(1412.29)=-2.388, p=0.017), and not for aDBS, (t(1412.24)=-0.776, p=0.438). There was no significant difference between the two interventions, (t(1412.04)=-1.620, p=0.105). No differences in errors were found between conditions, (Chi2(2)= 1.512, p=0.469).

Overall, stimulation was applied 48.8% (SEM 3.8) of the time during aDBS.

## Discussion

Our study provides the first pseudorandomised and blinded acute comparison of both UPDRS scores and side-effects in aDBS versus cDBS in the chronically implanted state. The results confirm previous findings that aDBS is well tolerated [9,11], performs equally well as cDBS, and does not compromise speech [14] in patients studied within days of electrode implantation. The effects in the chronically implanted state were seen despite stimulation being delivered about 50% of the time.

The patient cohort described in this study had DBS electrodes implanted for seven years on average (range 4-12 years), which provided the unique opportunity to observe the effects of aDBS in patients with advanced stages of the disease. It also confirms previous findings that beta oscillations remain informative of the clinical state of the patients [5,31], even several years after implantation. There was significant improvement of the blindly evaluated subUPDRS during aDBS and cDBS. A subanalysis of bradykinesia subscores showed that aDBS significantly reduced these items, but cDBS only showed a trend in this regard. aDBS and cDBS did not differ significantly and improvements across both groups were relatively modest. However, this might reflect the reduction in the response of bradykinesia items to stimulation reported in some of the studies following DBS patients for more than 5 years [3,4]. This may also explain why the level of improvement in subUPDRS scores with both aDBS and cDBS (∼20%) was at the lower limit of the range of improvements seen under similar blinded conditions when these interventions are contrasted acutely following electrode implantation [9,10,12,14,32]. Note too that scores from blinded video assessments are lower than those made during direct, unblinded clinical examination [9].

Regarding tremor subscores, both cDBS and aDBS significantly suppressed tremor, but cDBS tended to suppress tremor scores more than aDBS, although again the difference was not significant. In the five patients who presented tremor, tremor was suppressed in three of them during both aDBS and cDBS (Supplementary video 1), but two patients demonstrated re-emergence of tremor when stimulation was triggered off during aDBS (Supplementary video 2). This phenomenon was previously described in a similar proportion (2/12) of cases undergoing aDBS [18], and might be related to the fact that beta oscillations are only correlated with bradykinesia symptoms, but not tremor [5,6]. Although the majority of patients experience relief of tremor with aDBS, it remains unclear which factors determine when this is not the case. Advances in tremor detection using LFPs [33] may provide additional biomarkers to control tremor in patients in which tremor cannot be effectively suppressed using only beta-based algorithms. Meanwhile in the minority of patients in whom tremor breakthrough does occur during aDBS, trials should assess the efficacy of control algorithms that have a minimum which is a low, rather than zero, stimulation intensity. Alternatively, the period between triggered stimulation bouts could be reduced in the aDBS condition.

A large number of patients classified as tremulous PD patients in the clinical records were included in this study. This could be explained by our inclusion criteria, which required patients with chronically implanted electrodes who were cognitively preserved. It has been shown that tremor-dominant PD patients usually present a more favourable cognitive prognosis [34]. It has been observed that the presence of tremor in PD patients is not constant, and some patients who start with tremor as the main symptom, may switch to a bradykinetic phenotype as disease progresses [35].

The potential benefits of aDBS regarding dysarthria are promising. SID is one of the most common side effects of cDBS, with a prevalence of around 10% in treated patients [36]. This usually adds to speech problems which are already present as consequence of the disease [37]. Adaptive DBS may therefore particularly provide a useful alternative to cDBS in patients with SID.

We found non-significant differences between cDBS and aDBS in the tapping test. Iteration times were shorter with cDBS, even though bradykinesia UPDRS subscores were lower with aDBS. The contrast might be explained by the effects of break-through tremor in the minority of patients on aDBS. In the RST, an interference effect was found, but this was not affected by stimulation (either continuous or adaptive).

### Limitations

Some limitations of this study have already been highlighted, but others remain. Because of the intra-operative nature of this study, not all the UPDRS items could be assessed. As operations were performed under local anaesthesia, the time available to complete the protocol was limited. Therefore, the results may potentially be affected by incomplete washouts between stimulation conditions. Pseudo-randomisation should have helped limit the impact of incomplete washout and condition order was not found to be a significant factor upon analysis. Moreover, wash-out has been considered a two-step process, consisting of an initial fast decrease in stimulation’s therapeutic effect, followed by a further, slow decline. Our wash-out interval was sufficient for the former fast process [38,39]. Similar arguments may apply to the wash-in of effects, given that stimulation was only applied for about 5 minutes in each condition. The brief wash-out and wash-in periods necessitated by our intra-operative constraints should have served to reduce the differences between conditions, but significant differences were still detected. Another possible confound in the present methodology is that both aDBS and cDBS were limited to monopolar stimulation at either contact 1 or 2, and high voltages sometimes interfered with the recordings, which led to the use of submaximal voltages. The resultant stimulation choices may therefore have differed from the chronically optimised contact selection and stimulation voltage prior to battery change (see Table 1). Nevertheless, voltages, pulse width and stimulation frequency were kept fixed between the cDBS and aDBS conditions, and the optimal voltages were estimated based on the response to cDBS during the assessment.

Nevertheless, the selection of cDBS contacts was more constrained than usual and bipolar stimulation was avoided so as to keep the two stimulation conditions as similar to one another as possible. Future aDBS experiments using non-segmented or segmented octopolar DBS electrodes may allow for more versatile stimulation montages [40]. Finally, it is still possible that the effects demonstrated here do not persist in the chronic setting. Ultimately, chronic trials are necessary that contrast the best possible cDBS parameters established during chronic therapy with aDBS.

## Conclusion

The acute effects of beta-based aDBS are as effective as conventional DBS in chronically implanted patients, just as it is immediately after electrode implantation [9]. This is in line with the evidence that beta oscillations remain present and informative over time [41]. The present findings are important, as the assessments in chronically implanted patients are not confounded by microlesion effects. Furthermore, comparisons were performed with cDBS, using stimulation parameters that were optimised, with the exception of stimulation contacts. We found that SID was reduced with aDBS, most likely because stimulation was applied only as necessary. However, it remains to be seen whether aDBS remains effective with prolonged use, and whether stimulation algorithms have to be adjusted in some patients with tremor.

## Acknowledgements

We would like to thank Renske de Vries for the evaluation of the voice recordings, and Jasper Peeters for his assistance during the measurements,, and Alek Pogosyan for the facilitation of custom-made scripts.

---

Supplementary video 1: video recording of a tremulous PD patient with a good tremor suppression during both cDBS and aDBS

Supplementary video 2: video recording of a tremulous PD patient that presented re-emergent tremor during aDBS.

